# RBCeq: An Integrated Bioinformatics Algorithm Designed to Improve Blood Type Compatibility Testing

**DOI:** 10.1101/2021.01.13.426510

**Authors:** Sudhir Jadhao, Candice Davison, Eileen V. Roulis, Elizna M. Schoeman, Mayur Divate, Arvind Jaya Shankar, Simon Lee, Natalie M. Pecheniuk, David O Irving, Catherine A. Hyland, Robert L. Flower, Shivashankar H. Nagaraj

## Abstract

While blood transfusion is an essential cornerstone of hematological care, patients that require repetitive transfusion remain at persistent risk of alloimmunization due to the diversity of human blood group polymorphisms. Next-generation sequencing (NGS) is an effective means of identifying genotypic and phenotypic variations among the blood groups, while the accurate interpretation of such NGS data is currently hampered by a lack of accessibility to bioinformatics support. To address this unmet need, we have developed the RBCeq (https://www.rbceq.org/) platform, which consists of a novel bioinformatics algorithm coupled with a user-friendly web server capable of comprehensively delineating different blood group variants from genomics data with advanced visualization of results. The software profiles genomic data for 36 blood group systems, including two transcription factors and can identify small genetic alterations, including small indels and copy number variants. The RBCeq algorithm was validated on 403 samples which include 58 complex serology cases from Australian Red Cross LifeBlood, 100 samples from The MedSeq Project (phs000958) and a further 245 from Indigenous Australian participants. The final blood typing data from RBCeq was 99.83% concordant for 403 samples (85 different antigens in 21 blood group systems) with that listed from the International Society for Blood Transfusion database.

## Introduction

Transfusion has historically been an important facet of hematological care. Red blood cells (RBCs) are the most transfused blood product, with roughly 85 million RBC units being transfused per year to treat conditions ranging from severe anemia, sickle cell disease, severe hemorrhage, leukemia, or stem cell transplantation (1). While these transfusions are essential, patients requiring repetitive transfusion are at a high risk of alloimmunization that can lead to delayed or acute hemolytic transfusion reactions, fetal anemia, and complications during pregnancy (2). To comprehensively minimize alloimmunisation extended typing of other blood groups besides ABO, RH and K is a necessity. It includes the typing for over 320 antigens (1521 alleles) from other clinically significant blood groups (MNS, RH, LU, FY, JK, DI, DO, CO, H, XK, GLOB, and AUG (3-6) (7). Within these blood groups, antigens are population-specific, and are subject to varying rates of polymorphism, further complicating transfusion safety efforts (8,9). Clinical interpretation of genetic blood typing variants based upon the guidelines of the American College of Medical Genetics (ACMG) is recommended (10).

While effective for well-characterized and common variants, blood group typing conducted using traditional serological or molecular methods are alone insufficient for the characterization of blood group antigens that are rare, weakly detectable, recombinant (Rh, MNS), partial, or novel (11). Next-Generation sequencing (NGS) technologies can overcome the limitations of serological and SNP based molecular techniques and has the capability to characterize blood group gene variants in genetically diverse populations (12-14). Accurately predicting blood group phenotypes based upon NGS data requires immuno-genetic knowledge, given that multiple genotypes may result in the same phenotype (as with the ABO, MNS, LE, and XG groups), and not all blood group antigens are direct primary gene products (including the ABO, Rh, LE, and H groups). A number of different tools/algorithms implementing bespoke statistical and machine learning approaches have been designed to process NGS data (15,16). In order to achieve optimal sensitivity and specificity from these algorithms, a thorough understanding of the underlying informatics is required. The time spent analyzing and reporting blood group profiles depends on the complexity of a specific variant and on the skills of a given bioinformatician/geneticist (17). Without improvements to the bioinformatics underlying blood group genotyping, the costs associated with data processing, storage, and analysis will exceed the costs of the sequencing itself, thereby reducing the economic viability of such a strategy. As personal genome sequencing is forecast to become increasingly prevalent in the near future, there is a clear need for the optimization of the bioinformatics algorithms underlying sequencing and blood group characterization approaches.

Despite the clear value of applying NGS-based blood group characterization data to blood bank services, at present there is no single tool capable of facilitating complete and comprehensive automation of blood group characterization without any substantial computational and storage user requirements. Only two relevant tools are available at present: BOOGIE (18) and BloodTyper (19). The blood group prediction accuracy of the BOOGIE software is reported to be 94% for 34 blood groups in high-quality single nucleotide variants. Following analysis, this tool outputs the two most likely phenotypes for each blood group system in an individual along with a score, leaving the user to select the correct blood group phenotype. BloodTyper had a 99.2% accuracy for the prediction of blood group phenotypes but for only 12 blood group systems. Most importantly, both methods are command-line applications and requires the accessory of software with expertise in bioinformatics to install and run. One of the major theoretical advantages to NGS-based blood typing is the potential to discover novel antigens. The clinical utility of integrating NGS based blood typing in the Red Cell Reference Laboratory has been demonstrated to be beneficial at resolving complex serology problems arising from the novel or rare alleles altering or silencing blood group expression (17). Yet, neither of these extant software tools has such functionality nor leverages published population genomics information to annotate the frequency of detected antigen.

To efficiently analyze NGS data in the context of transfusion medicine applications, we have developed a novel algorithm and created a secure, comprehensive web server designated ‘RBCeq’. It can characterize not only known blood group alleles but also possible novel alleles capable of reducing and silencing antigen expression, functioning as web server-based blood group genotyping software that thus addresses both computational and storage challenges associated with the processing of large raw NGS datasets. Analyzing NGS data to predict blood groups is a complex and time-consuming task, and RBCeq addresses the unmet needs in this field and will facilitate the use and translation of this technology to improve blood donor and patient safety

## Materials and Methods

### Construction of a database of known blood group antigen alleles

A comprehensive blood group allele database was created using multiple manually curated data sources (International Society of Blood Transfusion (ISBT), Blood Group Antigen FactsBook(20), Human Blood Groups (21), Erythrogene (22), and RhesusBase (http://www.rhesusbase.info/). At the time of the design, 36 blood group system and 2 transcription factors (TF) were known, and the database is representative of the knowledge. The coordinates of known blood group antigen variants recorded in these databases were provided in the conventional cDNA reference sequence form. Corresponding blood group genotype coordinates for the hg19/GRCh37 genome were identified and validated using the Transvar (23), NCBI Clinical Remap (https://www.ncbi.nlm.nih.gov/genome/tools/remap), Ensembl Variant Effect Predictor (VEP) (24), and UCSC (25) resources. During the curation process, inaccuracies and omissions in the published antigen alleles (such as inconsistencies between reference nucleotide change and positions and amino acid identities and positions) were detected (Supplementary Table 1). These alleles were manually curated and validated using NCBI (https://www.ncbi.nlm.nih.gov/) and UCSC in order to ensure that the data were non-redundant and that allele names and corresponding phenotypes were uniform. The database was further improved using the previous publications (26,27) and extensive literature mining. In total, the resultant backend database is representative of approximately 1463 alleles from 44 genes, and two transcription factors (GATA binding protein 1: GATA1 and Kruppel like factor 1: KLF1) with 58 alleles, encoding blood groups arising from 36 blood group systems recognized by the ISBT.

### Variant calling

For variant calling, BAM files are processed in accordance with GATK4 best practices (28) which includes first pre-processing with BaseRecalibrator, ApplyBQSR, and then variant calling using HaplotypeCaller. Variant calling and haplotype phasing are conducted for a restricted set of 44 blood group genes and 2 transcription factors (GATA1 and KLF1)(Supplementary Table 2).

### Characterization of clinically relevant and novel blood group alleles

The filtered high-quality variants that are not mapped to known blood group alleles are queried against the ClinVar database (29). The mapped variants are reported as clinically significant variants, while the remaining variants are processed for rare variant analysis. RBCeq checks the frequency of the variant in the gnomAD database (∼143,000 genomes). If the frequency is less than 0.05 (or less than a user-defined threshold) and the respective variant is nonsynonymous or a splice-site variant, then that variant will be reported as a rare. Remaining variants that are not clinically relevant or rare will be processed for novel variant analysis. To filter novel SNVs, RBCeq uses six independent computational tools (SIFT, Polyphen2, MutationTaster2, LRT, FATHMM, and PROVEAN) to assess the impact of genetic variants on protein structure and function. If any one of these tools determines the variant to be deleterious, and it is nonsynonymous or a splice-site variant, then the variant will be reported as a novel (Figure 1).

**Figure 1:**
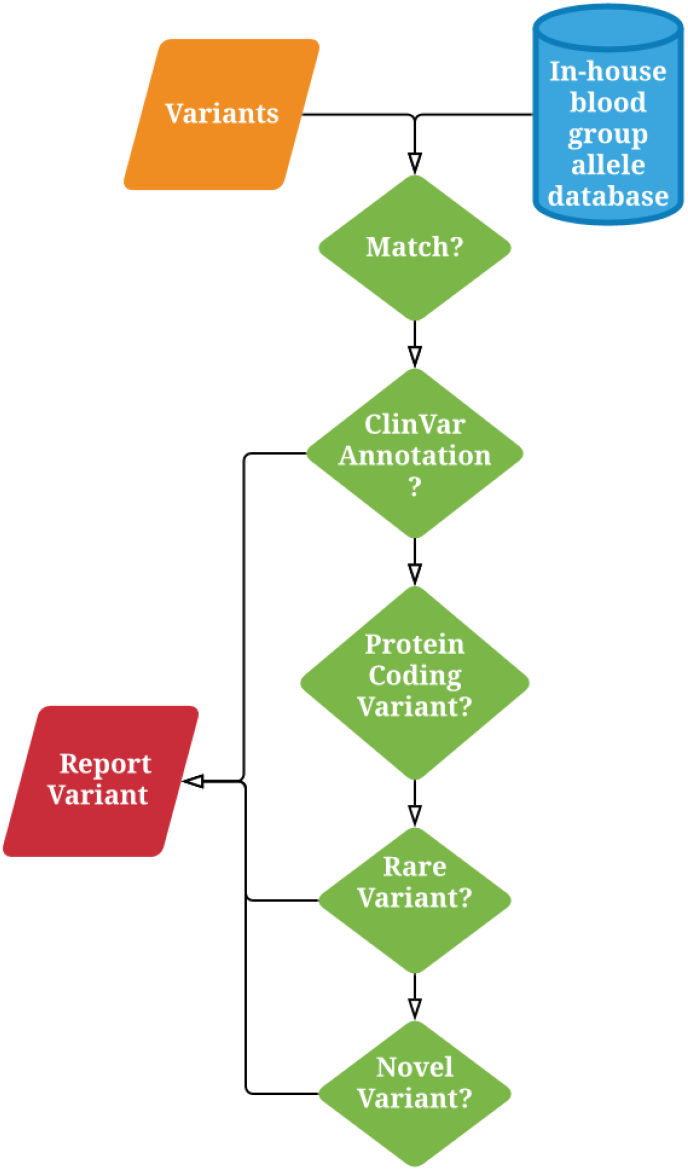
Workflow for the annotation of clinically significant, rare, and novel blood group variants.

### Datasets

To evaluate the performance of RBCeq, we accessed the following datasets.

#### Proof of Principle Algorithm Development dataset

Initial development was undertaken using 13 in-house targeted exome sequencing (TES) samples for which serological data for 14 blood groups (+ 4 samples with SNParray) were available with the remaining 22 blood groups were predicted(27), provided in the supplementary table 3. In addition 45 previously published complex blood group serology cases with TES, from the Australian Red Cross Lifeblood Red Cell Reference Laboratory with manually predicted blood groups (Supplementary Table 4) (17,27,30) were analysed.

#### South East Queensland Indigenous TES data

245 targeted blood group exome sequencing samples with serological phenotypes for ABO, D, C, c, E, e, K and k blood groups(30). This dataset was important to validate the accuracy of the algorithm with established methods currently used to determine ABO, D, C, c, E, e, K, k phenotypes. No other data sets provided this amount of serological data.

#### MedSeq project

110 whole-genome sequencing samples (30X) from the MedSeq Project randomized controlled trial (accession number phs000958) were accessed through dbGaP authorized access.

#### The 1000 genomes (1000G) project

The 1000G project includes 2504 whole-genome sequencing (WGS) samples from 26 population groups classified into 5 super populations. 2504 WGS bam files were accessed through the 1000G project FTP server.

#### Erythrogene

Erythrogene is a database of the predicted blood group genotype information for 2504 WGS samples from the 1000G Project. The blood group genotype information was accessed through www.erythrogene.com.

#### gnomAD

gnomAD is the aggregation of 125,748 exome sequences and 15,708 whole-genome sequences from unrelated individuals sequenced as part of various disease-specific and population genetic studies. The data is available through the Broad Institute FTP (https://gnomad.broadinstitute.org/downloads), and consists of data from the following populations - AFR: African/African American, AMR: admixed Americans, ASJ: Ashkenazi Jewish, EAS: East Asian, FIN: Finnish, NFE: Non-Finnish Europeans, OTH: Other population, and SAS: South Asians.

## Results

### Creation of blood group genotype and phenotype prediction algorithm

The genotype/phenotype prediction process is composed of three stages: optimal allele selection, dominant/recessive pair selection and allele pairing (*AP*) score calculation. During optimal allele selection, each variant (*V*) is scanned against all blood group (*bg*) allele (*A*) variants for an exact match. The blood group alleles (*A*_*bg*_) with at least a single variant match are then selected.

### Optimal allele selection

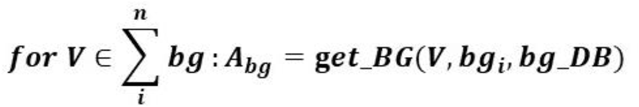

**KEY: V: Variants; bg: blood groups; A: All possible Alleles; bg_DB: blood group allele database**

During dominant/recessive pair selection, every selected optimal allele will be paired (*a*_*1*_, *a*_*2*_) against each other such allele. Alleles with homozygous identical genotypes or heterozygous differing genotypes will be paired (compatible (*a*_*1*_, *a*_*2*_)). An AP score (*S*_*a1,2*_) is then assigned to each allele pair to select a pair with a high propensity to affect the phenotype of the impacted gene. The allele pair with the lowest AP score (*min*_*APS(S)*_) will be used during the final genotype and phenotype calling. If the algorithm fails to find an alternate allele for a particular blood group, then the reference genotype and phenotype will be reported.

### Dominant/recessive pair selection

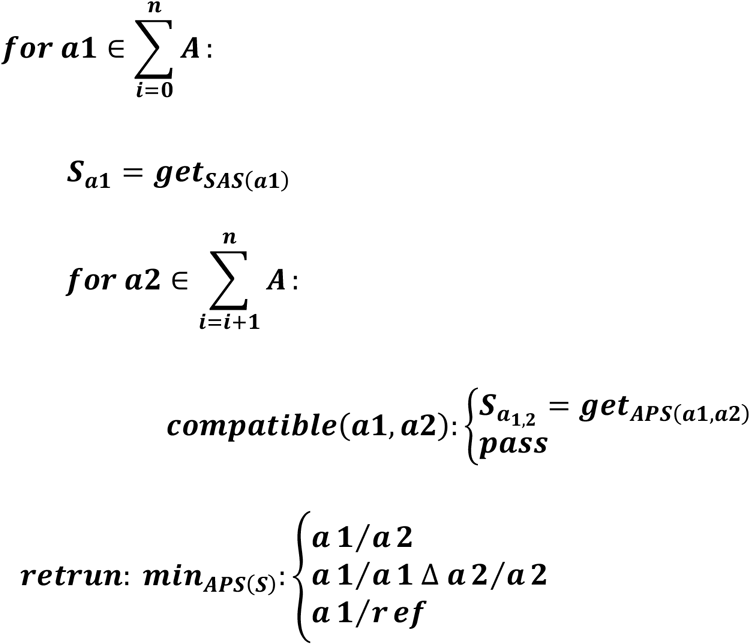

**Key: SAS: Single allele score, APS: Allele pair score, S: Allele pair, ref: Reference alleles**

### Allele pair score (APS) calculation

AP scoring was devised as a novel approach to prioritizing optimal allele pairs with the potential to alter gene phenotypes. These scores were specifically designed to address genotype/phenotype predictions in complex blood group systems (e.g., ABO, Rh, MNS, and Lewis) where certain haplotypes share some identical variants in between them, making it possible to detect more than two allele pairs in a given sample. These allele pairs may have a complete or near-complete correspondence to known allele profiles. In such scenarios, the allele pair with the highest number of variants matching the genotype will be prioritized by the scoring system even if an allele with fewer variants exhibits complete correspondence (Figure 2).

**Figure 2:**
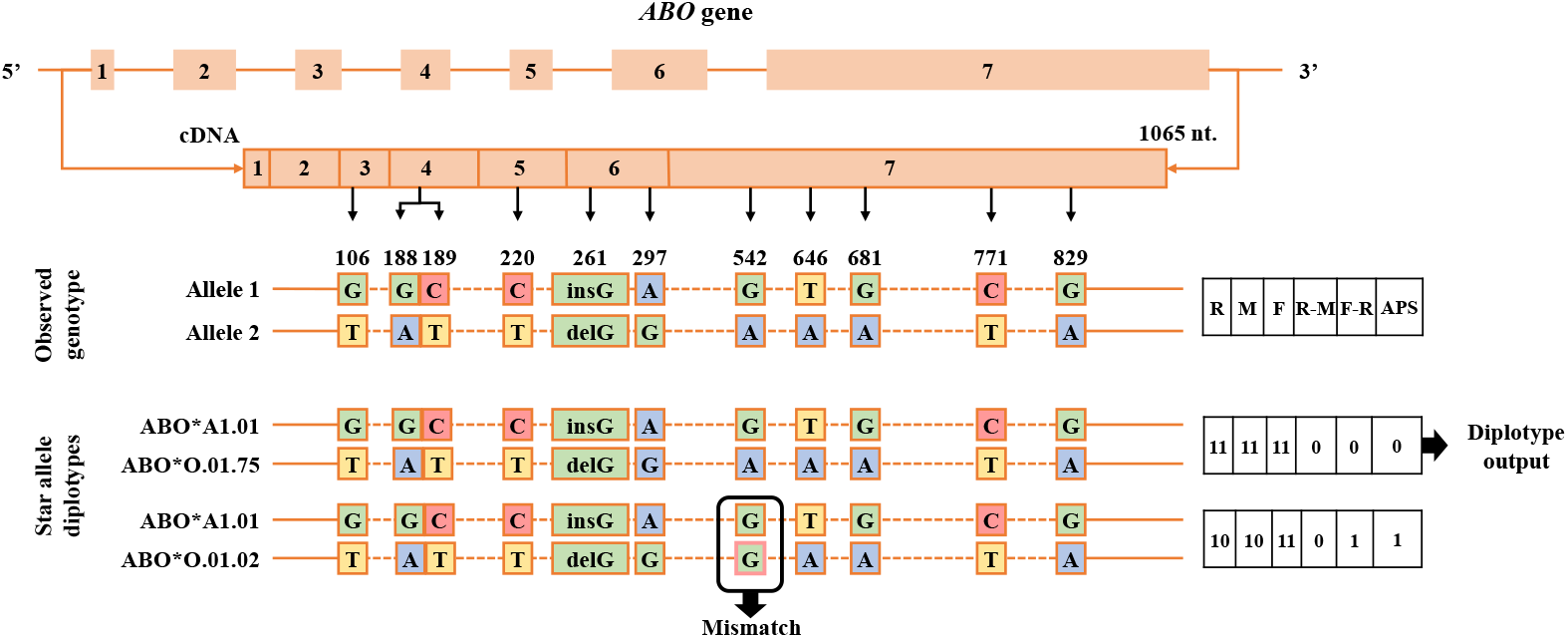
The breakdown of APS calculation and selection of best allele pair to predict ABO genotype for NA21112 sample from 1000G dataset. The c. 542G>A variant associated with ABO blood group present in sample genotype but not needed for the definition of the allele ABO*0.01.02.

### APS calculation

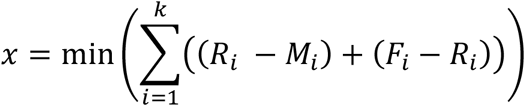

**Key: R: the total number of variants required to define a blood group allele; M: the number of variants matched; F: the total number of variants found in the sample file for that blood group; K: the total number of alleles**.

First the number of variants (R) required to define an allele pair is calculated (*ABO*\**A1*.*01/ABO*\**O*.*01*.*75*: 11; *ABO*\**A1*.*01/ABO*\**O*.*01*.*02*: 10) then how many are concordant (M) with input sample genotype (*ABO*\**A1*.*01/ABO*\**O*.*01*.*75*: 11; *ABO*\**A1*.*01/ABO*\**O*.*01*.*02*: 10), and finally the total (F: 11) ISBT or RBCeq database known variants for that particular blood group system is calculated. The subtraction R-M, are the variants which are missing in the input sample genotype but required in the definition of the associated allele. In this case, both allele pair have complete correspondence, so R-M is zero for both. The subtraction of F-R, are the variants present in the sample genotype but not required in the definition of associated allele pair. The allele pair *ABO*\**A1*.*01/ABO*\**O*.*01*.*75* (F-R: 0) is defined by 11 variants and all of them are present in sample genotype whereas in the definition of *ABO*\**A1*.*01/ABO*\**O*.*01*.*02* allele pair c. 542G>A is not required (F-R: 1). The addition of R-M and F-R values for each allele pair indicates the number of variants present in the sample genotype not associated with the allele pair call (*ABO*\**A1*.*01/ABO*\**O*.*01*.*75*:0 *ABO*\**A1*.*01/ABO*\**O*.*01*.*02*: 1). Therefore the allele pair call with the lesser number means the allele pair chosen makes the most out of the observed genotype (*ABO*\**A1*.*01/ABO*\**O*.*01*.*75*: 0) and vice versa (Figure 2).

### Algorithm Validation

The RBCeq algorithm was validated on 403 samples which include 58 complex serology cases from Australian Red Cross LifeBlood, 100 samples from The MedSeq Project (phs000958) and a further 245 from Indigenous Australian participants. The algorithm was initially iteratively developed and validated on 13 samples for which serological data were available for 14 blood groups (+ 4 samples with SNParray) and remaining 22 blood groups with manual prediction provided (Supplementary Table 3). Cis-trans haplotype ambiguities were solved in most cases using the allele pair scoring system. We observed that certain haplotypes as defined in the ISBT database, did not require all variants as part of the allele definition to be present for a particular phenotype (e.g. *ABO*\**O*.*01*.*02*: c.829G>A). In this case, if partial allele matches are found, only nonsynonymous variants and their zygosity alone was used to determine the most likely genotype and phenotype. Previously published CNV based genotyping approach for the RH blood group system were adopted in RBCeq (19,31,32), enabling the correct definition of the RHD, C and c antigens on 13 training samples (Supplementary Table 3). We further validated the algorithm on 45 complex blood group serology cases from the Australian Red Cross Lifeblood Red Cell Reference Laboratory (Supplementary Table 4) with predicted phenotype. The initial iteration of the algorithm predicted two discordant results, where it missed calling *ABO*\**O*.*02*.*01* allele which is associated with c.802G>A instead of c.261delG.

Next, the low frequency alleles (e.g., *GYPB*\**06*.*01, LU*\**02*.*19, KEL*\**02*.*10*) of MNS, LU and KEL blood groups were detected at the heterozygous level, but the algorithm missed predicting the low-frequency antigen. The algorithm was further improved to identify the heterozygous low frequency antigens phenotype change and the O antigen genotyping with respect to both O alleles (*ABO*\**O*.*02*.*01* and *ABO*\**O*.*01*.*01*).

Additionally, our algorithm was validated on The MedSeq Project (phs000958) comprising of 100 whole-genome sequencing and respective serology samples, and it achieved 100% accuracy across all the blood groups (Supplementary Table 5). To further develop and qualify the accuracy of RBCeq as a means of reporting blood groups through comparisons with serology results, we analyzed serological blood group data (ABO, Rh) for 245 Indigenous Australian individuals to those predicted using RBCeq, again achieving 100% accuracy for ABO blood group profiles. In two samples, RBCeq predicted discordant calls for C antigen, which was due to inaccurate read mapping in the homologous region of exon2 RHCE.

In summary, our validation approach successfully detected 85 antigens in 21 blood group systems (Supplementary Table 6). However, the NGS based blood group phenotyping is highly dependent on data coverage and quality of input sequence data, and our algorithm allows user-supplied coverage and quality cut-off as input. As such, users must exercise caution when defining run parameters and interpreting reported blood group profiles. The collective accuracy for 21 blood group systems in 405 samples was 99.83%, with discordant results only for RHCE

### Webserver

#### Implementation

We developed a webserver to enable the seamless analysis of blood group profiles from NGS data. The web server was developed using Apache and PHP and is hosted on Amazon Web Services (AWS), thus making it scalable. The interactive visualizations are implemented through d3.js and c3.js libraries. RBCeq uses EC2 container service-Docker management on AWS to serve several different concurrent users/jobs. The primary advantage of RBCeq is that it is personalized for every user such that each user must first create a login in order to use it. The user-based login allows users to obtain blood group profiles independently and seamlessly in parallel with many simultaneous sessions using an organized queue-based system. It also enables users to save their outputs and to access their project at a later date or from a different place, enabling user mobility and collaboration. User data uploaded into the server is only utilized for RBCeq analyses, and will be stored for six months, after which it will be erased. Users can also delete their data sooner or export the data from RBCeq in a user-friendly Excel spreadsheet containing run parameters, dates, times, input file names, uploaded data types, and complete blood group profiles for archiving purposes. Thorough website penetration testing and social engineering of the RBCeq webserver has been undertaken to assess for vulnerabilities and potential security risks associated with environment and the technology used in building server and deemed secure according to OWASP (Open Web Application Security Project) guideline (https://owasp.org/www-project-top-ten/).

### Web Interface-Input Files and Analysis Component

Once the user has logged in, interactions associated with job submission are facilitated by the “create job” tab which follows a relatively straightforward stepwise process based on the format of the input files for processing. The user has the option to upload a BAM file, or VCF and BAM files and define the run parameters for RBCeq file processing to support consistent and reproducible variant calling outputs (Allele Depth, Genotype Quality, MAF) (Figure 3). To overcome large BAM file size and associated uploading issues, we have developed a stand-alone GUI tool named BAMTrimmer that can be downloaded directly from RBCeq and can run in Windows operating systems. The BAMTrimmer removes all unmapped and duplicate reads and trims the files with respect to blood group-associated genes. It thereby reduces BAM file size significantly, allowing users to upload these files to RBCeq within a few minutes. Once the trimmed BAM and compressed VCF (vcf.gz) are uploaded, the user can define run parameters and submit the job for processing. The uploaded file will be validated to ensure they adhere to their respective format guidelines.

**Figure 3:**
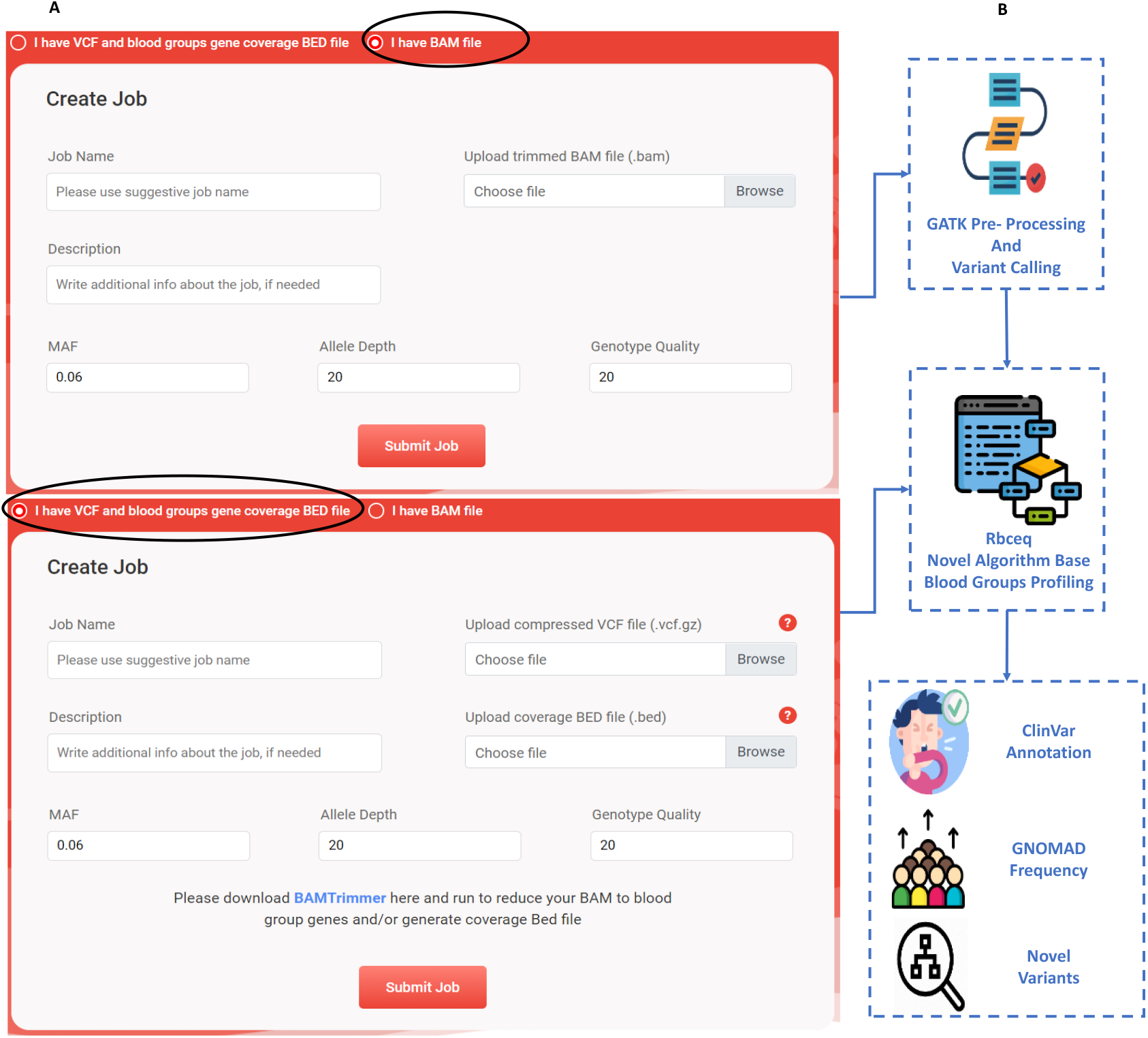
A. A screenshot illustrating the web interface used to upload different Input file formats and the parameters required to run RBCeq. B: The workflow of analysis component of RBCeq

The RBCeq analysis component is compartmentalized into three sequential parts: 1. Variant calling, 2. Known blood group profiling, and 3. Annotation of non-ISBT variants with respect to clinical significance, population frequency distributions, and variant novelty. BAM files will be processed in accordance with GATK4 (28) best practices as previously described. [32] The RBCeq server determines the blood group profiles according to novel decision algorithms developed in-house (Figure 3).

### Result Visualization

RBCeq was developed with a specific emphasis on multiple approaches to visualize data by representing data using both interactive graphics and dynamic tables. Once jobs are completed the output can be visualized by the user by the “View Output” tab, which will direct the user to the result page. Details pertaining to each results section are given below:

### Job Summary

This section describes information pertaining to user-specified input file format and run parameters used, including input file job name, and RBCeq unique job ID number, run time and date, and provides an option to download completed results as an Excel file (Figure 4).

**Figure 4:**
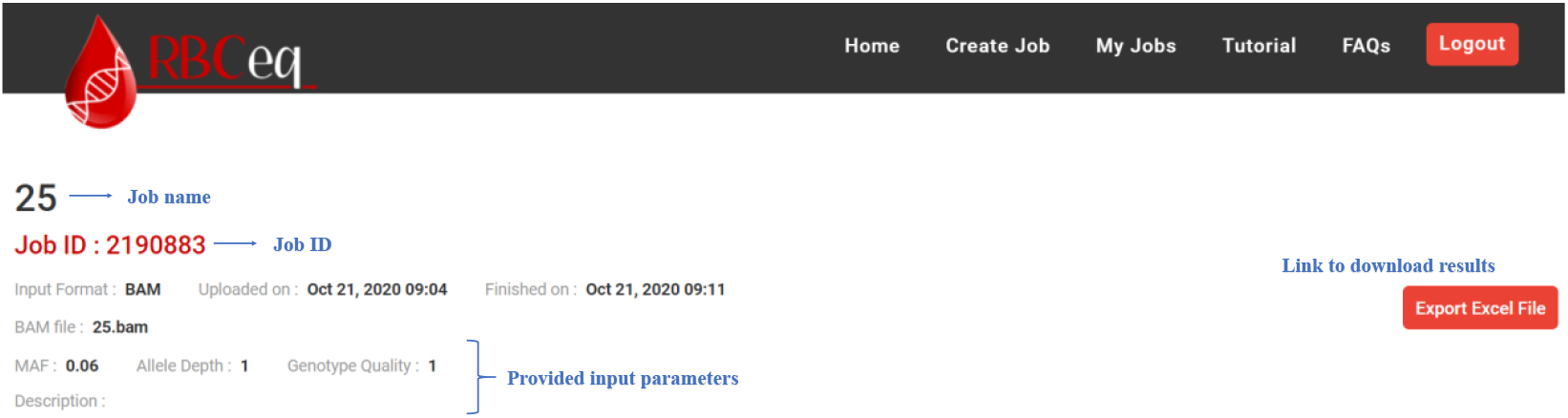
Screenshots illustrates the overview of the job summary.

### Overall Summary

This section includes a summary of variant annotation and average sequence read mapped coverage in blood group associated genes. The interactive pie chart serves as a visual reference for variant annotation distributions. The overall summary provides a quantitative overview of user-provided files, enabling a rapid bioinformatics quality check of inputted samples (Figure 5A).

**Figure 5A:**
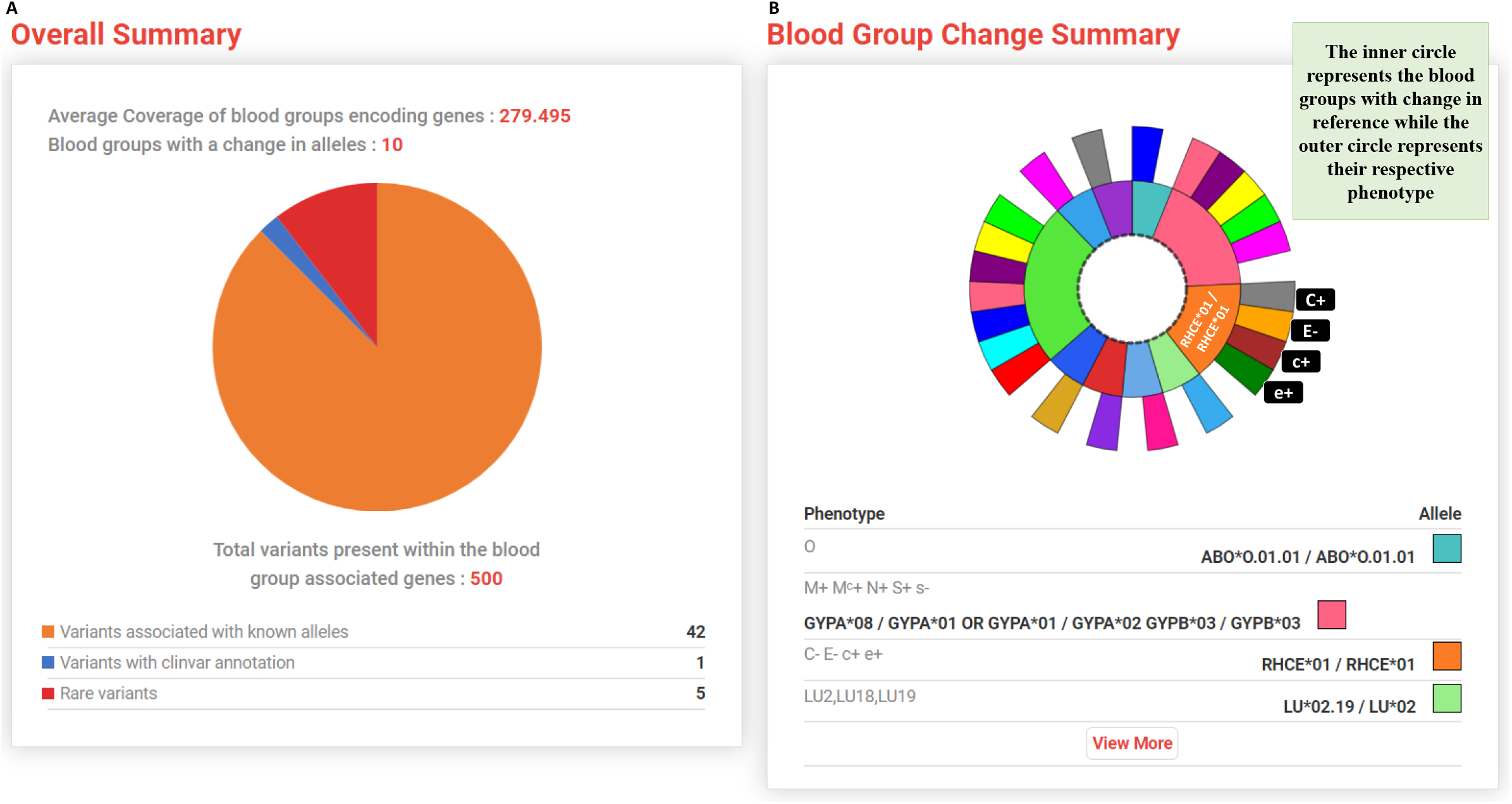
The summary of distribution and annotation of detected variants in the blood group antigen defining genes; includes average coverage of blood group defining genes; the number of blood groups alleles for which the reference is changed, and the distribution of total variants detected in blood group associated genes with blood group known, ClinVar and rare annotation. B: The interactive blood group change allele pie chart.

### Blood Group Change Summary

This section summarizes the detected known blood groups alleles that are different from the reference together with information regarding the associated phenotype. Colours in the inner circle represent the blood group alleles that are changed relative to the reference, while the colours of the outer circle represent respective phenotypes. Users can interactively explore genotype and phenotype information pertaining to detected blood groups by moving their cursor on the plot (Figure 5B).

### Per Blood Group Variants and Coverage Statistics

This section provides an interactive graph that enables the quantitative analysis of gene coverage and detected variants for each blood group. The plots will help the user visualize correlations between read mapped coverage and detected variants for all blood group genes. Users can also zoom in or out on a particular gene by scrolling their mouse wheel over the plot (Figure 6A). Figure 6B highlights the table of variants detected for each blood group gene.

**Figure 6:**
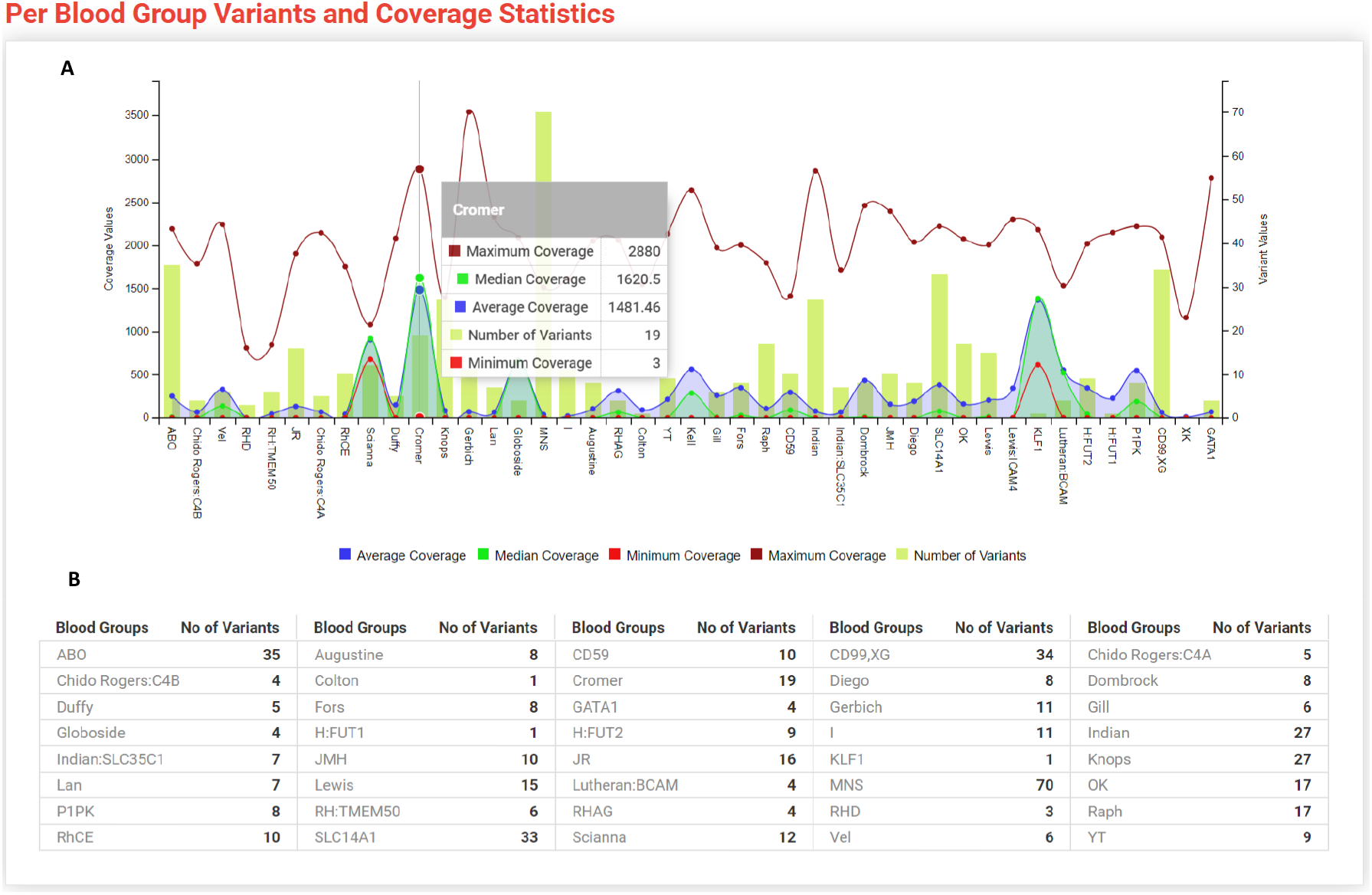
The “Per Blood Group Variants and Coverage Statistics” graph is an interactive graph which gives the quantitative analysis with respect to blood group genes coverage and detected variants. A: Line/bar combo plot, the x-axis is the blood group antigen defining genes, left y-axis represents the coverage of the genes, and the right y-axis represents the number of variants for each blood group antigen determining genes. Each line colour represents different coverage value. B: The table gives the number of variants detected in the input sample for each blood group antigen defining genes.

### RH Blood Group System Coverage Statistics

This section describes the CNV calculator and predicted results for phenotyping/genotyping based on analysis coverage statistics extracted from the uploaded trimmed BAM file for the RH (*RHD* and *RHCE* genes) blood group system known for structural variation between the homologous genes. The CNV ratios and interpretation for exonic rearrangements or deletion/duplication/triplication are given (Figure 7).

**Figure 7:**
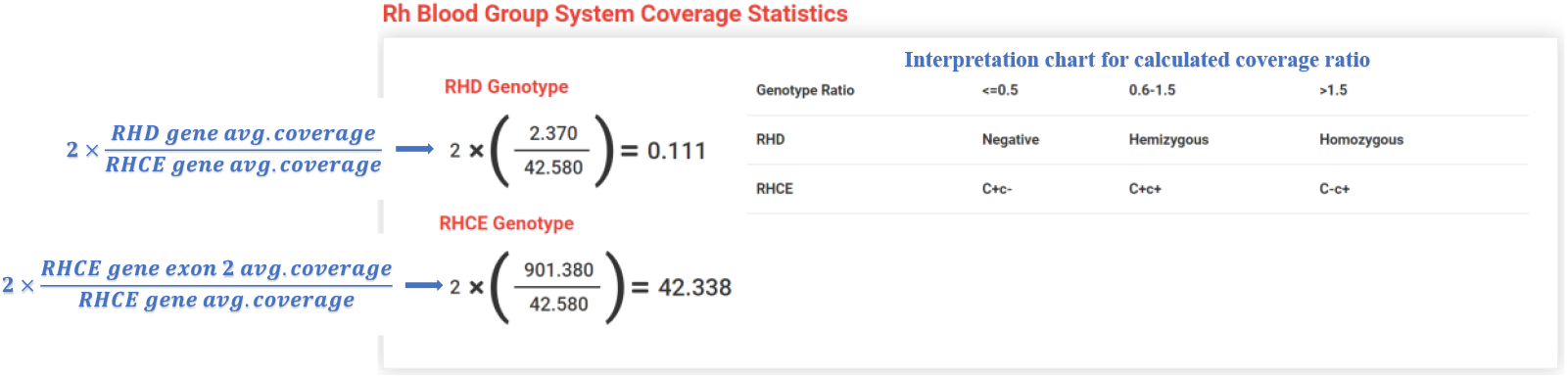
The CNV ratio formula (RHD=2* (*RHD* gene average Coverage/*RHCE* gene average Coverage and RHCE = 2* (*RHCE* gene exon2 average coverage/*RHCE* gene average coverage) and interpretation for exonic rearrangements or deletions/duplications/triplications.

### The Known Blood Group Allele table

Reference and blood group alleles that are different from the reference source will be reported along with supporting information including variants, zygosity, allele depth, and allele frequency (Figure 8).

**Figure 8:**
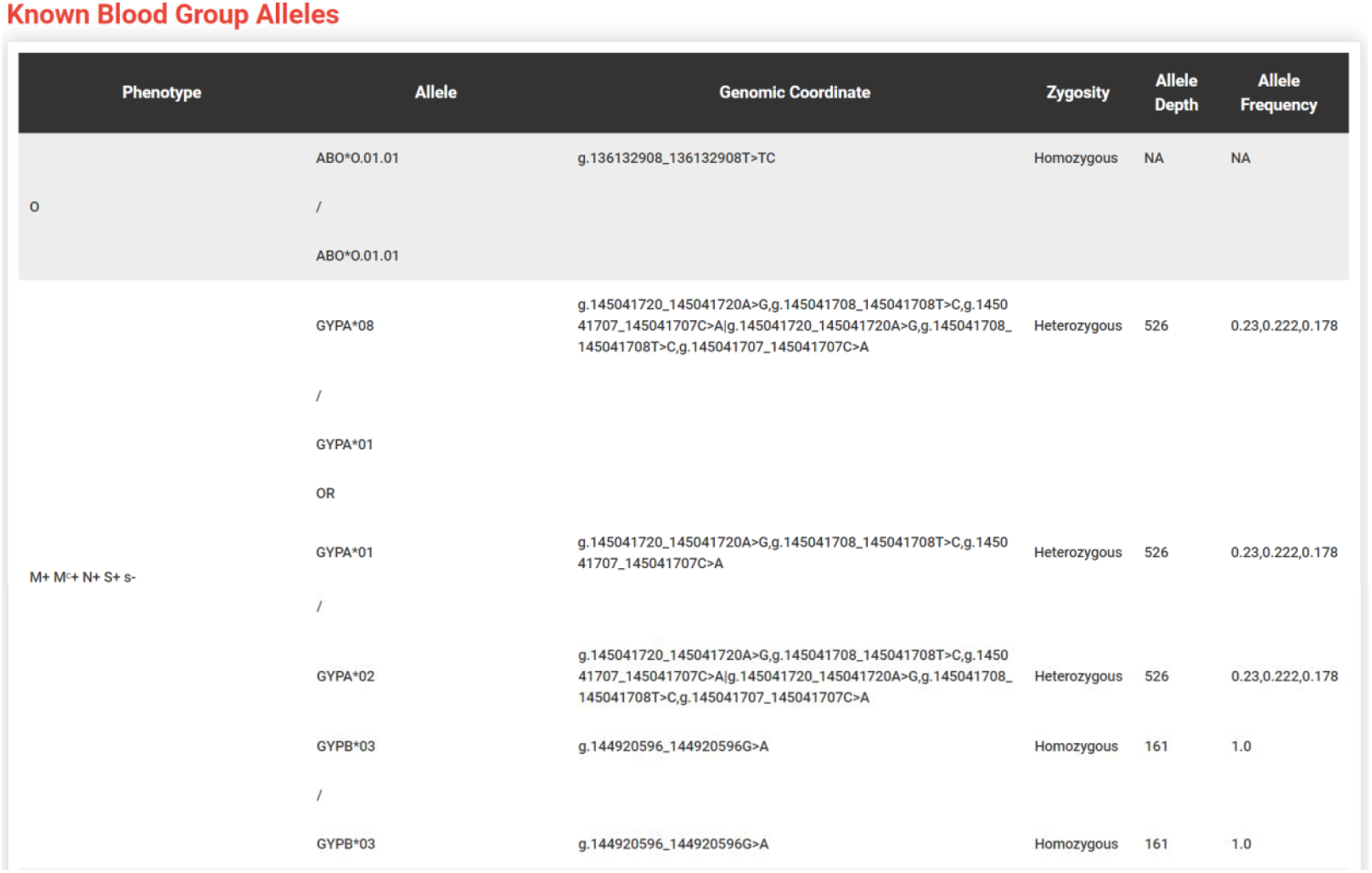
An excerpt of the Known Blood Group Allele table; The first column provides details pertaining to the blood group phenotype, while the second column includes allele information, the third column contains information regarding the variants responsible for defining the allele, the fourth column includes the zygosity of the allele (if one variant is heterozygous then the whole allele will be referred to as a heterozygous allele), the fifth column includes the average allele depth for that allele, and the sixth column includes the average allele frequency. For the complete table of 36 blood group profile with 2 TF, please follow the tutorial page at https://www.rbceq.org/tutorial.php.

### Clinically Significant, Rare, and Potentially Novel Allele Annotations

This section partitions and lists the variants into 3 categories (Figure 9); according to the algorithm/workflow described in Figure 1: ClinVar, Rare, and Novel variants.

**Figure 9:**
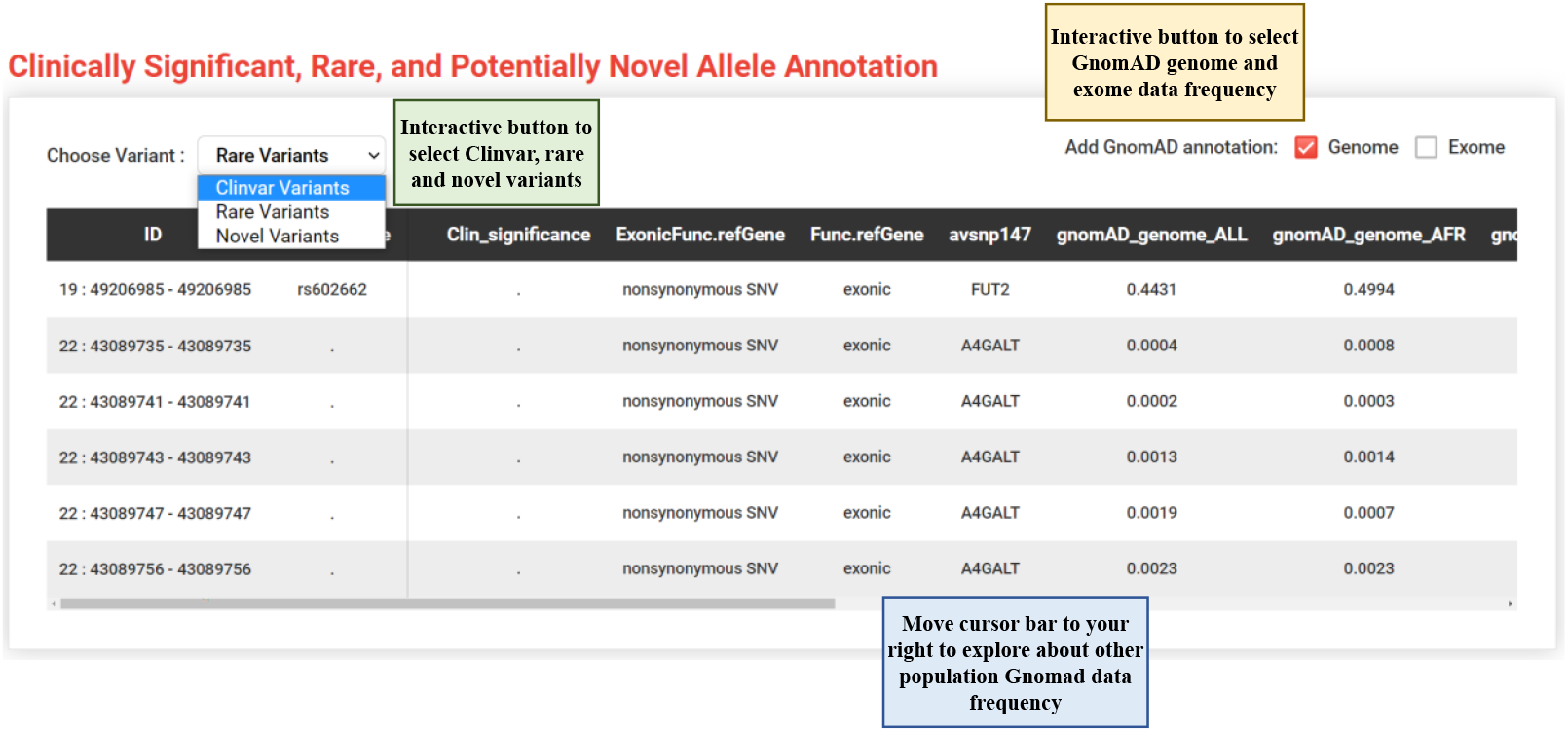
The first option includes ClinVar Variants, which are variants that have ClinVar annotations. The second option includes Rare Variants that have a MAF less than the user-selected threshold value in the gnomAD database. The third option includes novel variants, that are predicted to be deleterious using *in silico* tools and with exonic function that is predicted to be “nonsynonymous / frameshift /stop-gain/stop-loss/splicing”. All lists are provided with known details pertaining to each variant, including dbsnpid, exonic function, refGene and gnomAD genome and exome frequencies in 9 different populations.

### Case Study 1: Extensive blood group characterization in diverse populations

In order to evaluate the capability of RBCeq to comprehensively characterize the blood group genotypic landscape of diverse populations, we obtained blood group genotypes from the gnomAD dataset (33) which lacks associated sample-level information or phased genotypes. Despite this limitation, it remains a valuable resource when assessing variant data and the prevalence of particular variants in different population. A minimum allele frequency approach was used to call multi-variant blood group alleles, such that all variants defining a given blood group allele needed to be present within a particular population at a frequency greater than zero.

A total of approximately 180,000 variants were detected across blood group encoding genes, of which only 1,107 have previously been reported to define blood group alleles. RBCeq predicted 805 (51%) non-reference blood group alleles out of 1521 known such alleles using a minimum allele frequency approach. Of the 805 identified ISBT alleles in gnomAD, 30% (474) were considered rare (<=0.05 MAF) in gnomAD genome, and another 41% (652) were rare in the gnomAD exome dataset. The further distribution of the total number of non-reference blood group alleles (805) was assessed to identify polymorphisms, with the ABO (17%), RHD (24%), RHCE (9%), KEL (5%), LE (5%), and H (6%) blood groups being among the most polymorphic in this analysis (Supplementary Table 7). Most of these alleles and their variants have a frequency of less than 0.3 across all eight populations. However, we also detected population-specific alleles that were clinically significant, such as *FUT2*\**01W*.*02*.*01*, which was present in ∼40% of EAS samples and considered rare in all other populations. Conversely, ABO*AW.25 is rare in EAS samples but common in other populations. Overall, 60 alleles were found to be completely absent in EAS populations that were relatively common in other populations. The RHD*01W.33 allele is common in NFE (non-Finnish European) populations but not in other populations, while the *ABO*\**O*.*01*.*09, ABO*\**AW*.*09, RHD*\**38, RHD*\**03*.*08, RHCE*\**02*.*30, RHCE*\**01*.*20*.*01, RHCE*\**01*.*20*.*02, JK*\**01W*.*03, JK*\**01W*.*04, KN*\**01*.*07, KN*\**01*.*06*, and *ABCB6*\**01W*.*02* alleles are common in African populations but rare in all other populations. Interestingly, the *FY*\**01N*.*01* allele which is protective against *P. vivax* infections in regions where malaria is endemic, was found to be common in the African population with a MAF 0.8 and rare in all other populations (Figure 10) (34). An additional 144 alleles that exhibited uneven frequency distributions across these populations were also identified (Supplementary Table 8). This analysis has revealed the breadth of blood group diversity over several world populations that can be extracted efficiently by RBCeq from WGS and whole exome sequencing (WES) data. This information is key to understanding population-level diversity, and would offer considerable benefits in supporting the integration of NGS in blood donor testing in the population.

**Figure 10:**
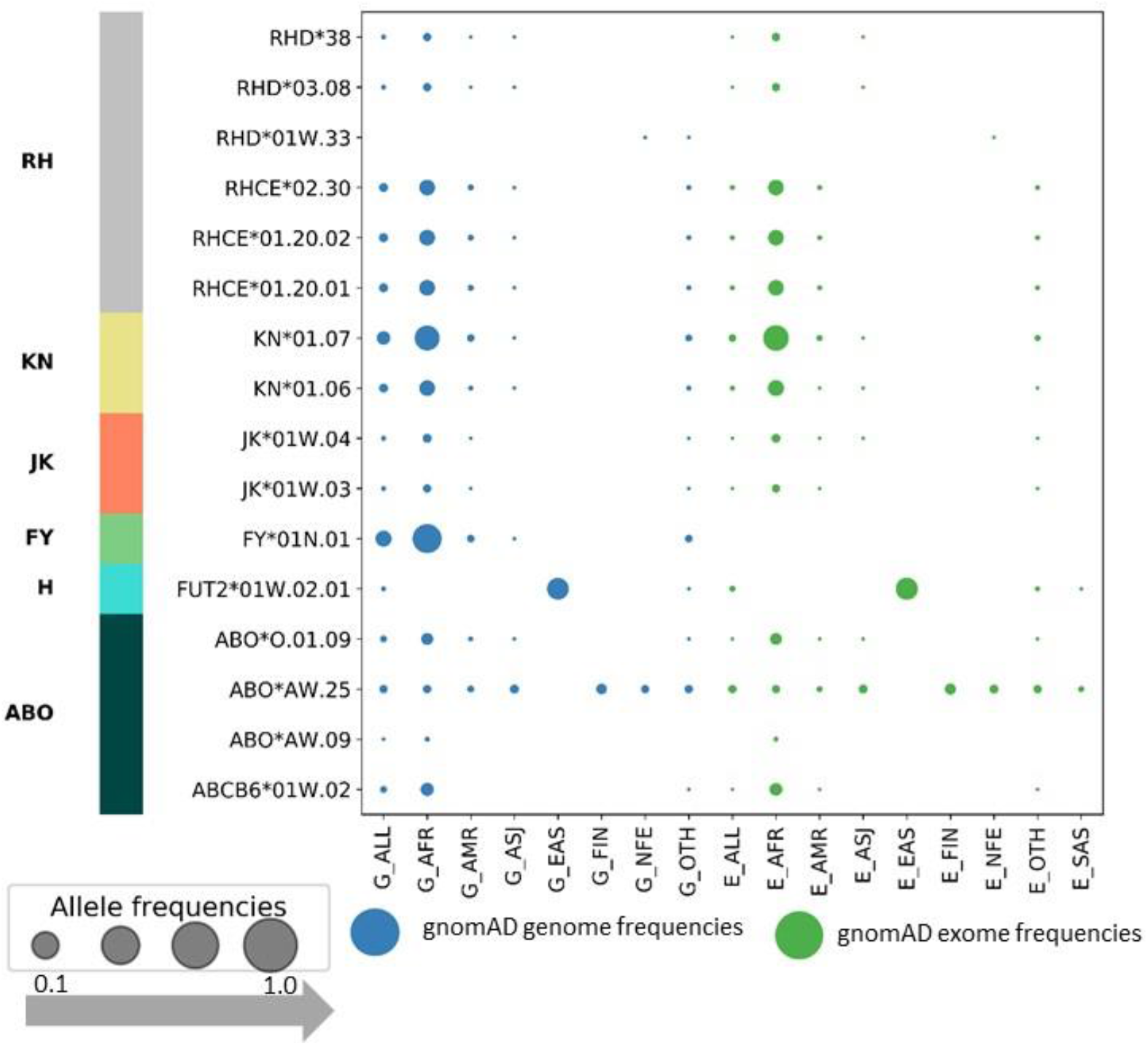
The population-specific alleles which are also clinically significant with MAF from gnomAD data. The frequencies in blue colour are from 15,708 WGS and green colour from 125,748 exome samples.

In a secondary analysis, RBCeq was used to characterize variants that had not previously been reported to be associated with blood group alleles but that had the potential to affect the antigen structure formation. We detected 52 (Supplementary table 9) such variants that also had clinical associations, ∼16,160 rare variants with frequencies of ≤ 0.0005 in any of the eight populations, and, most importantly, we identified 159 (Supplementary table 10) variants that were computationally predicted to be novel and deleterious (Figure 11). Interestingly, no novel variants associated with the genetically complex ABO blood group system were identified, and as little as six novel alleles associated with the RH blood group system. This observation is expected from these two most commonly studied systems. These numbers prompt further scientific enquiry to better understand the clinical significance of these variants arising from other blood group systems. As such, rapid and comprehensive characterization of blood group genetics is possible when analyzing NGS data using RBCeq or similar tools, thereby providing more information on blood groups profiles which can be used clinically to minimize the risk of transfusion-related complications.

**Figure 11:**
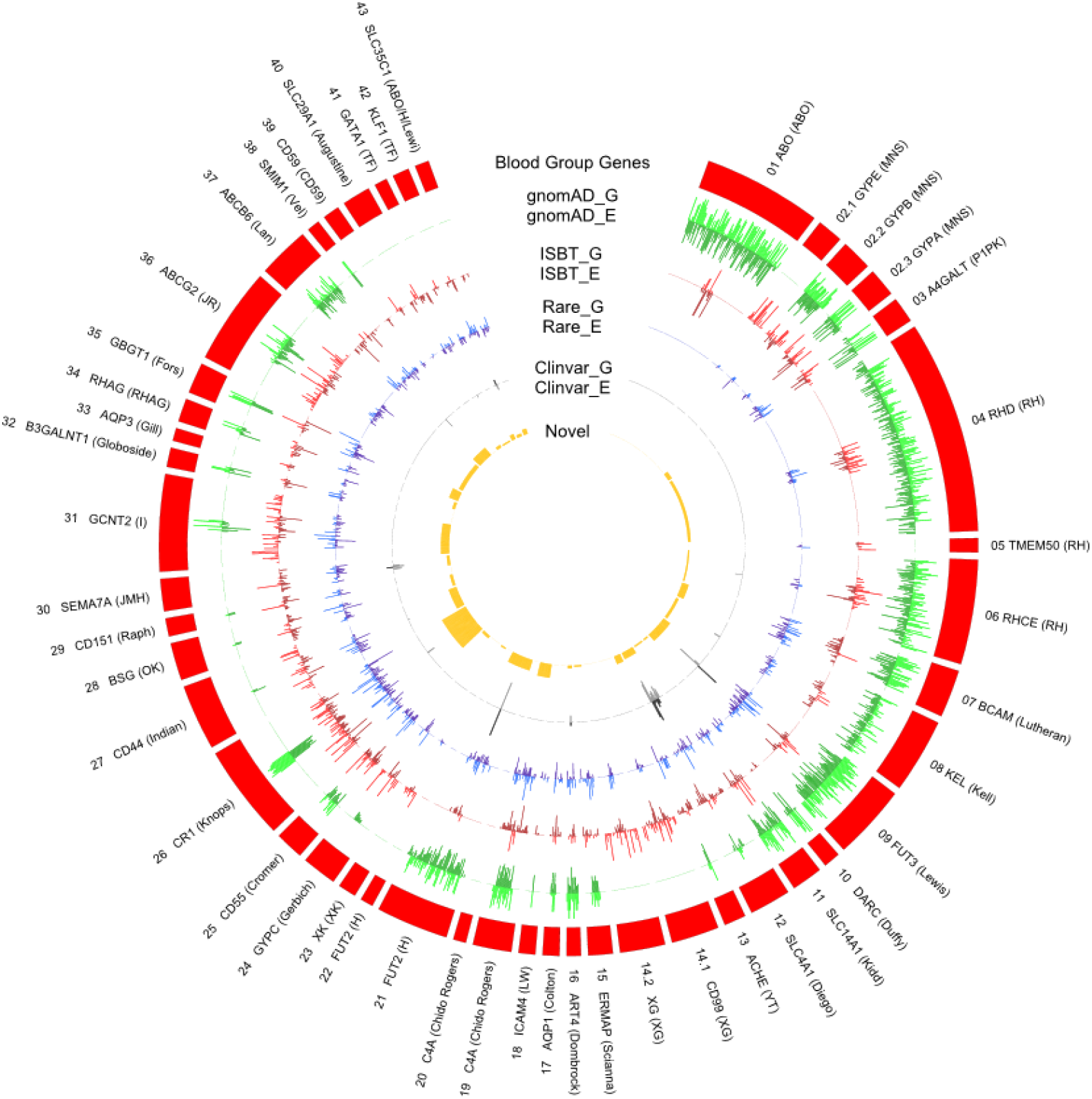
The distribution of gnomAD (genome and exome) datasets genetic variants and their frequency in RBC antigen encoding genes. The outer ring (red) represents the RBC antigen encoding genes; box length represents the number of variants observed; G is denotes gnomAD genome frequency and E is denotes gnomAD exome frequency. The outer green (light/dark) circle indicates the distribution of variants frequency across different blood group genes from the gnomAD data. The red (light/dark) circle indicates the number of variants with ISBT associations relative to all gnomAD variants. The blue (light/dark) circle indicates the distribution of rare non-ISBT variants in all six populations. The dark grey circle indicates the number of non-ISBT variants annotated to the ClinVar database. The yellow circle shows the distribution of number of novel variants.

### Case Study 2: 1000G-based blood group profiling

To further demonstrate the processing power and utility of RBCeq as a means of analyzing diverse WGS data pertaining to 5 super population groups (AFR, AMR, EAS, EUR, SAS), we predicted the blood group profiles associated with the 1000G dataset (2504 samples) (35). The blood group genotype profiles corresponding to the 1000G data have previously been published by Moller et al. (22). A comparison at the allele frequency level was not possible because allele frequency assignments in Erythrogene are made with respect to extra novel alleles that are not reported in the ISBT blood group allele database, and the sample-level blood group profile is not given by the Erythrogene database. RBCeq predicted 74 more alleles than reported by the Erythrogene database and has assigned revised comprehensive genotypes to the 1000 genome dataset (Supplementary Table 11). Overall, the Erythrogene database contains ∼3340 alleles, of which 255 are known to the ISBT with a frequency greater than zero in the 1000G samples. Of these 255 alleles, seven were missed by RBCeq calling - the variants that define the *KEL*\**02M*.*04, JK*\**01N*.*09, RHD*\**01EL*.*36, RHD*\**01W*.*28*, and *RHD*\**59* alleles were not present in any of the VCF files for the 1000G dataset. The ISBT allele designation for the *AUG*\**02* allele is NM_001304463: c.1171G>A, whereas in the Erythrogene database it is associated with NM_001304463: c.1297G>A. With respect to ABO allele predictions, *ABO*\**O*.*01*.*58* allele variants were present in a large number of samples and were checked in a reduced sample set where we found that our scoring system outperformed the Erythrogene genotype predictions. For example, in sample NA21137, a total of 20 (c.106G>T, c.188G>A, c.189C>T, c.220C>T, c.261delG, c.297A>G, c.488C>T, c.526C>G, c.53G>T, c.595C>T, c.646T>A, c.657C>T, c.681G>A, c.703G>A, c.771C>T, c.796C>A, c.802G>A, c.803G>C, c.829G>A, c.930G>A) known variants were present that were associated with the ABO blood group (NM_020469.2), and all are covered in the *ABO*\**O*.*01*.*68* allele, whereas *ABO*\**O*.*01*.*58* is defined by just seven of these variants (c.261delG, c.297A>G, c.646T>A, c.681G>A, c.687C>T, c.771C>T, c.829G>A). As such, the *ABO*\**O*.*01*.*58* allele did not maximally utilize the available variant data, whereas RBCeq selected the *ABO*\**O*.*01*.*68* allele that utilized 9 present variants from the VCF file. RBCeq detected 74 different alleles that were not reported in Erythrogene. The 74 alleles are associated with the ABO, MNS, RH, LU, KEL, FY, JK, YT, H, and KN blood group systems. The details of each allele identified are provided in Supplementary Table 11. We were unable to ascertain why these alleles were not reported in Erythrogene, but it may have been affected by their algorithm having also defined new alleles based on genotype. The Erythrogene study was published in 2016, after which many new alleles have been reported in the ISBT database.

We also detected 11 ClinVar (Supplementary table 12), and 1093 rare (Supplementary table 13) non-ISBT SNVs in 1000G datasets. RBCeq used an average of 1.20 minutes and 15 Mb of memory to predict a blood group profile on each sample of 1000G from a VCF file. This case study demonstrates the processing power and capability of RBCeq to manage analyzing large WGS datasets and providing additional insights into the blood group profiling of the extensively researched 1000 genome dataset. All 2504 sample blood group profile results for this case study are available on the RBCeq website (https://www.rbceq.org/).

### Comparison with Existing Tools

In contrast to other tools, RBCeq is user-friendly, fast, accurate, and provides extended profiles of variants with no current known blood group phenotype association (Table 1). BOOGIE (18) and bloodTyper (19) do not have the capability of data pre-processing for variant calling, despite being the necessary first step prior to variant calling. One of the significant theoretical advantages to NGS-based blood typing is the potential to discover rare and novel antigens, yet neither of the extant software tools has such functionality. RBCeq annotates blood group gene variants with clinical significance (ClinVar), rare and novel variant with gnomAD frequency. The BOOGIE tool reports an accuracy of 94% for 34 blood groups. BloodTyper reports the accuracy of 99.2% but has been evaluated for only 21 antigens from 12 blood group systems (14 blood group genes). In contrast, the RBCeq evaluation included 85 different blood group antigens from 21 blood group systems (21 genes). In comparison to other tools, RBCeq is readily accessible and available online to all users for research and academic purposes. RBCeq is also efficient with processing time and memory, permitting streamlined analysis, blood group profile report and interactive visualisation of sample data with an average of 5 million reads (BAM) in 7 minutes. The detailed feature base comparisons of RBCeq with published tools are given in Table 1. The RBCeq integration provides user-friendly open-source software that will boost the adoption of blood group detection and prediction using NGS data.

**Table 1:**
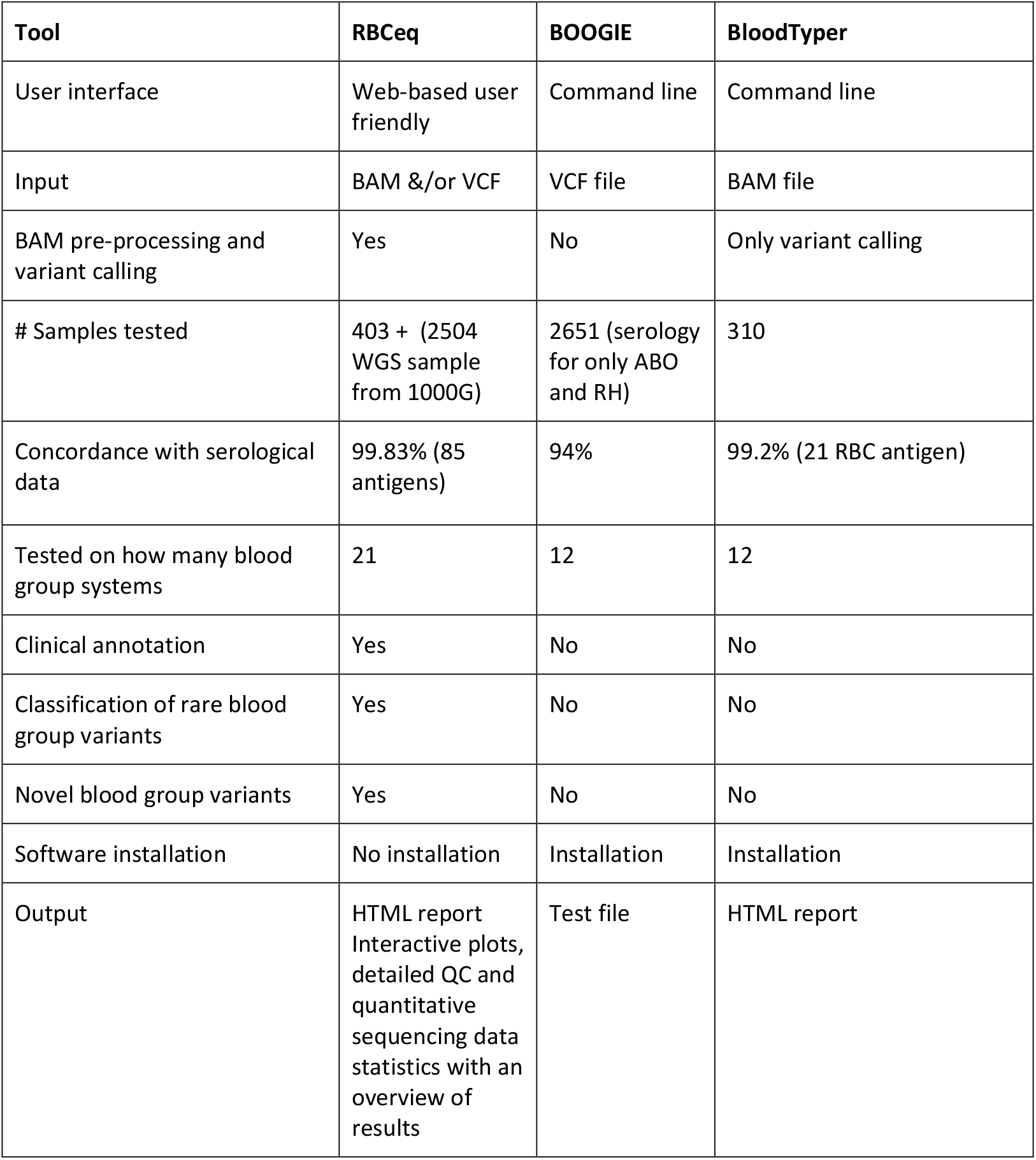
Comparison of the features of RBCeq to those of existing tools

| Tool | RBCeq | BOOGIE | BloodTyper |
| --- | --- | --- | --- |
| User interface | Web-based user friendly | Command line | Command line |
| Input | BAM &/or VCF | VCF file | BAM file |
| BAM pre-processing and variant calling | Yes | No | Only variant calling |
| # Samples tested | 403 + (2504 WGS sample from 1000G) | 2651 (serology for only ABO and RH) | 310 |
| Concordance with serological data | 99.83% (85 antigens) | 94% | 99.2% (21 RBC antigen) |
| Tested on how many blood group systems | 21 | 12 | 12 |
| Clinical annotation | Yes | No | No |
| Classification of rare blood group variants | Yes | No | No |
| Novel blood group variants | Yes | No | No |
| Software installation | No installation | Installation | Installation |
| Output | HTML report<br>Interactive plots, detailed QC and quantitative sequencing data statistics with an overview of results | Test file | HTML report |

## Discussion

RBCeq is the first scalable web-based blood profiling platform to be implemented online, enabling users to define analysis parameters, produce iterative interactive visualizations, and to archive their results. The collective accuracy for 21 blood group systems in 405 samples is 99.83% with discordant results only for RHCE. The user can effectively analyze blood group profiles corresponding to 36 blood groups and two TFs based upon WGS, WES, or TES data, extracting non-ISBT variant information to gain a more profound understanding of the underlying data.

Previously published tools such as BOOGIE were developed to analyze just 34 blood group systems, and validation was constructed for just the ABO and D blood groups with a serological concordance of 94%. The bloodTyper tool achieved a concordance of 99.2% for 21 antigens in 12 blood group systems (14 blood group genes) when used to analyze 310 samples. In contrast, our RBCeq algorithm achieved a concordance of 99.83% for 85 antigens in 21 blood group systems across 403 samples. The subtypes and co-dominant antigens of the ABO blood group were not validated by bloodTyper, whereas we were able to validate ABO subtypes (A1, A2, B, and B3) and A1B co-dominant antigens. RBCeq also managed to accurately report complex and rare blood group phenotypes observed in a Red Cell Reference setting such as Hy-, which weakens DO blood group expression and is rare in the African populations, as well as in Vel-, which is present in 2.56% and 0.6% of Scandinavian and African populations, and Jk(a+^w^b-), which is common in most populations(36). Integrating NGS-based blood typing in the Red Cell Reference Laboratory represents a clinically beneficial approach to resolving the complex serology problems that arise from novel or rare alleles which alter or silence blood group expression, yet neither of the extant software tools has such functionality (37,38).

Previously overlooked non-ISBT variant information has the potential to be integrated into clinical decision systems in order to support transfusion efforts in individuals that do not harbor common antigens in an effort to reduce the risk of antibody sensitization associated with conditions including Hemolytic Disease of the Fetus and Newborn (HDN), severe anemia, leukemia, sickle cell disease. For example, one prior complex HDN analysis from a collaborative laboratory revealed that secondary analyses of NGS data were able to detect novel low-frequency antigens present in < 1% of the global population, including the AUG:3/ATML+ and SARA+/MNS:47 antigen. Extensive serological testing for these antigens failed to reveal any antibody specificity in these patients, and the reagents necessary to detect these rare/novel blood types are available in only a small number of reference laboratories in the world. Cases of this type require rapid antibody identification for optimal management of suspected feto-maternal incompatibility and risk of bleeding at delivery (37,38).

In future releases, we plan to incorporate a platelet antigen prediction function. Furthermore, in the time since work commenced on the development of the RBCeq platform, five new blood group systems have been recognized by the International Society of Blood Transfusion Red Cell Immunogenetics and Blood Group Terminology Working Party (ISBT 037 KANNO, ISBT 038 SID, ISBT 039 CTL2, ISBT 040 PEL and ISBT 041 MAM). These blood group systems will be included in future releases of RBCeq. In the context of the automation of hybrid alleles (e.g. RHCE*ce-D(4-7)-ce, GPB(1-46)-A(47-118)) prediction is highly dependent on statistical normalization and batch-corrections methodologies. To automate the hybrid allele detection process, far clearer descriptions of the methods are required to facilitate robust predictions. Even so, as the number of relevant studies continues to increase, these complex phenotypes may be better clarified (39-41).

RBCeq holds great promise for use in the automation of blood group trait detection in personalized medicine applications, given the constantly rising number of identified phenotypes, potentially aiding global blood supply organizations in the computational genotyping of all clinically relevant blood groups for their large numbers of their blood donors. In so doing, RBCeq will help to ensure transfusion safety for all populations, supporting equitable access to quality healthcare.

## Acknowledgement

We acknowledge and pay respects to the First Nations Peoples of Australia who were part of the South East Queensland Indigenous study to develop & validate RBCeq to predict their unique blood group profiles accurately. We are grateful for the opportunity to work together and the trust placed in us to undertake research in a respectful and collaborative manner. This study was conducted in collaboration with the Carbal Medical Services in Toowoomba (https://carbal.com.au/), in consultation with the Indigenous Community Advisory Committee and under the approval of the Australian Red Cross Lifeblood Human Research Ethics Committee (2018#17 /QUT ethics approval IDnumber). The authors also thank Maree Perry, Aoibhe Mulcahy and Glenda Millard for their role in sample receipt, preparation, serological testing and sequencing (TES) of the Indigenous samples.

